# Xylan inhibition of cellulase binding and processivity observed at single-molecule resolution

**DOI:** 10.1101/2024.01.29.577775

**Authors:** Nerya Zexer, Alec Paradiso, Daguan Nong, Zachary K. Haviland, William O. Hancock, Charles T. Anderson

## Abstract

Efficient cellulose degradation by cellulase enzymes is crucial for using lignocellulosic biomass in bioenergy production. In the cell wall of plants, cellulose is bound by lignin and hemicellulose, which are key factors contributing to the recalcitrance of plant biomass. These non-cellulosic cell wall components are known to interfere with the function of cellulolytic enzymes. While the effects of lignin have been studied extensively, the contribution of xylan, the major hemicellulose in the secondary cell walls of plants, is often overlooked. To study those effects, we generated model cell wall composites by growing bacterial cellulose supplemented with varying concentrations of purified xylan. We used single-molecule microscopy to image and track fluorescently labeled *Tr*Cel7A, a commonly used model cellulase, as it binds and hydrolyses cellulose in these synthetic composites. We found that minute amounts of xylan are sufficient to significantly inhibit the binding of Cel7A to cellulose. The inclusion of xylan also reduced considerably the proportion of moving enzyme molecules, without affecting their velocity and run length. We suggest that, when available at low concentrations, xylan thinly coats cellulose fibrils, and incorporates as continuous patches when available at higher concentrations. Non-productive binding of Cel7A to xylan was not found to be a major inhibition mechanism. Our results highlight the importance of targeting xylan removal during biomass processing and demonstrate the potential of using single-molecule imagining to study the activity and limitations of cellulolytic enzymes.

**Graphical abstract:** 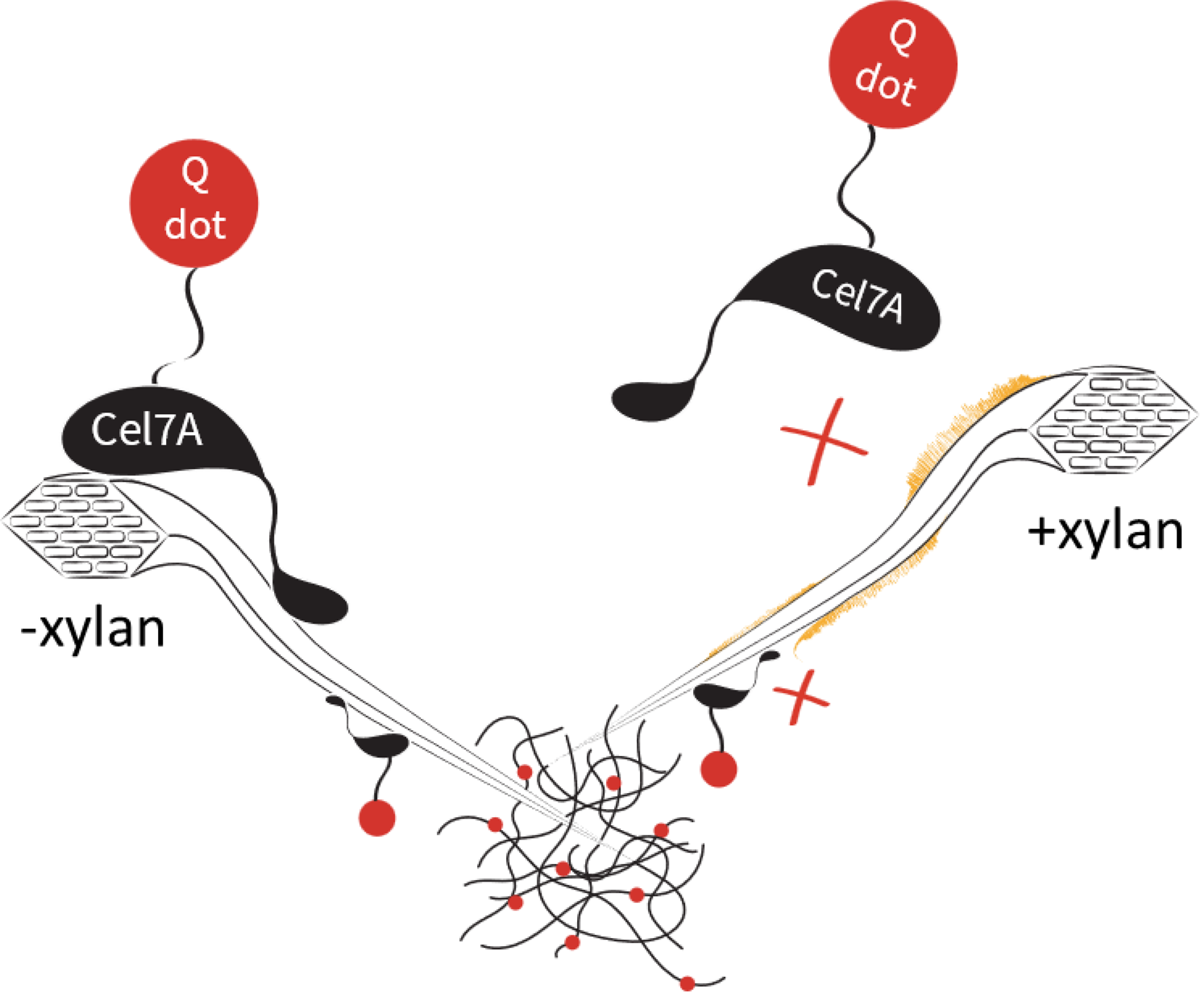

## Introduction

The secondary cell walls (SCW) of plants are complex composites composed of cellulose, hemicellulosic polysaccharides, lignin, and proteins, and are crucial for water transport and mechanical support in plants. They also represent the most significant renewable resource on the planet. This so-called lignocellulosic biomass is the major raw material for several industries including papermaking and construction, and in the last two decades has been the focus of bioenergy production.

Composed of a β-1,4 xylose backbone, xylan is the most abundant hemicellulose in plant SCW. It coats and crosslinks bundles of cellulose microfibrils, and is proposed to function in part as a transition phase between the rigid cellulose fibrils and non-crystalline polysaccharides and lignin^1,2^. Chemical modifications of the xylan backbone and its degree of polymerization vary between species and cell types. Xylan modifications can include glucuronic acid, galacturonic acid, rhamnose, and acetyl groups. In eudicot SCW, highly acetylated glucuronoxylan is the dominant form. The pattern and degree of these modifications influence how xylan interacts with itself and other cell wall components^2,3^. The genes involved in xylan biosynthesis are known and some aspects of biosynthetic processes are well-characterized, with polymerization and decoration occurring in the Golgi apparatus and secretion to the apoplast occurring concurrently with SCW cellulose deposition^4^.

The depolymerization of cellulose during biofuel production depends on the activity of cellulase enzymes. One commonly used model cellulase is *Tr*Cel7A, a cellobiohydrolase from *Trichoderma reesei* (teleomorph *Hypocrea jecorina*). This exo-cellulase degrades cellulose by engaging with the reducing ends of glucose chains and hydrolyzing glycosidic bonds to release cellobiose. This process, assisted by a carbohydrate-binding module (CBM), requires the extraction of a single glucose chain from the cellulose lattice, and its threading into the enzyme’s tunnel-shaped catalytic domain (CD). The enzyme remains complexed with the cellulose as it processively advances along the fibrils^5,6^. Previous works showed that these processive runs can terminate by dissociation of the enzyme-glucose complex, but slow de-complexation that leads to stalled enzyme molecule that remains unproductively engaged with the substrate^7^ had also been suggested as the rate-limiting step^6,8^.

Efficient enzymatic saccharification requires adequate access of cellulase enzymes to the cellulose substrate. The abundance of xylan in plant biomass and its close contact with cellulose make this hemicellulose a potential determinant of the recalcitrance of plant biomass. Xylan is hypothesized to hinder cellulase access to its substrate, and supplementation of cellulases with xylanases increases glucose conversion from pretreated corn stover and poplar biomass^9,10^. Xylan has also been suggested to increase biomass recalcitrance by blocking the catalytic site of cellulases and competitivity inhibiting these enzymes^11,12^. However, because work to date has relied on bulk biochemical assays, the specific molecular mechanisms by which cellulase activity is inhibited by xylans are not well understood.

Several species of gram-negative bacteria synthesize cellulosic biofilms^13^, a highly pure form of cellulose that has many applications in medical and food industries. Understanding its unique physiochemical characteristics is of interest for developing bio-based composites and polymers research^14,15^. Bacterial cellulose is similar in its chemical structure to plant-derived cellulose, but tends to form larger fibrils, and is highly pure as it does not contain lignin, pectin, or hemicellulose, making it an attractive substrate for cellulase activity assays^16,17^.

To directly examine how xylan interferes with cellulose degradation by Cel7A, we produced composites made of bacterial cellulose and purified xylan by synthesizing the cellulose in the presence of varying concentrations of beechwood xylan. We used immunofluorescence and monosaccharide analysis to validate xylan incorporation, and interference reflection microscopy (IRM) and scanning electron microscopy (SEM) to structurally characterize the cellulose-xylan composites. Using total internal reflection fluorescence microscopy (TIRFM) to image quantum dot (Qdot)-labeled Cel7A, we tracked and analyzed the dynamics of individual enzymes as they bound to and degraded the synthetic composites. We show how small amounts of xylan can inhibit Cel7A binding and motility on cellulose, with implications for bioenergy production strategies.

### Experimental

#### Preparation of composites

Primary cultures of 50 mL Schramm-Hestrin (SH) media were inoculated with a fresh colony of *Gluconacetobacter hansenii* (ATCC 23769), and incubated at 30°C while shaking at ∼180 rpm for 48 h. Secondary cultures were prepared in sterilized two-liter glass trays by adding 500 mL of SH media with appropriate amounts of beechwood xylan (Megazyme) and 50 mL of the primary culture. Trays were covered with aluminum foil and incubated uninterrupted for five days at 30°C until cellulosic pellicles had formed.

#### Substrate processing

Harvested pellicles were washed three times with dH_2_O followed by a 70% ethanol wash and three more washes of dH_2_O, then transferred to 0.5M sodium hydroxide and incubated at 80°C for 30 minutes, and again washed with dH_2_O. To neutralize the pH following sodium hydroxide incubations, pellicles were washed three times with sodium acetate buffer (50 mM, pH 5.0), followed by three more dH_2_O washes, and stored at 4°C. Using scissors, pellicles were shredded into small pieces, suspended in 30 mL dH_2_O, and sonicated with a Sonic Dismembrator (Thermo Fisher, model 100) five times for 30 seconds each (intensity setting 9), at one-minute intervals. Sonicated cellulose suspensions were further processed with a microfluidizer (Microfluidics, model LM20). Samples were run through a 100 µm chamber (Microfluidics, H10Z) at 15,000 psi for 10 minutes (∼20 cycles for 30 mL samples).

#### Immunofluorescence labeling and staining

For immunofluorescence labeling of xylan, 10 µL of composite suspensions were pipetted on glass microscope slides and dried at 40°C for two hours. Dried samples were encircled using a liquid blocker PAP pen (Electron Microscopy Sciences) to contain labeling and washing solutions. Samples were blocked for 30 min with phosphate-buffered saline (PBS) containing 1% w/v bovine serum albumin (BSA). After blocking, samples were washed with PBS and incubated overnight in blocking solution with 1:10 diluted LM11 anti-heteroxylan antibody (PlantProbes) at room temperature. The next day, samples were washed three times with PBS, and incubated with Alexa 488 goat-anti-rat IgM secondary antibody (Jackson ImmunoResearch) diluted 1:100 in blocking solution for 90 min at room temperature in the dark. To stain for cellulose, samples were rinsed three times with PBS, and stained for 10 min using 0.01% (w/v) S4B (Pontamine Fast Scarlet 4B, marketed as Direct Red 23 by Sigma, catalog # 212490).

#### Fluorescence imaging and colocalization analysis

Xylan immunofluorescence labeling and cellulose S4B staining of samples were imaged on a Zeiss Cell Observer SD microscope with a Yokogawa CSU-X1 spinning disk unit. Images were collected using 40X and 100X oil objectives. A 488 nm excitation laser and a 525/50 nm emission filter were used to image the Alexa Fluor 488 signal, and a 561 nm excitation laser and a 617/73 nm emission filter were used to image the S4B signal.

Colocalization of Alexa 488 and S4B signals was quantified using the ImageJ PSC colocalization plugin ^18^. Analysis was performed on at least six images collected using a 100X objective from two experimental replicates for each substrate. The entire field of view was used for the analysis and the plugin threshold value was set to zero.

#### Composite hydrolysis and monosaccharide analysis

To prepare samples for monosaccharide analysis, 2 mL of each composite suspension was centrifuged at 20,000 x g for five minutes. The supernatant was discarded and the collected substrates (∼10 mg) were subjected to acid hydrolysis using 1 mL of 72% sulfuric acid for 5 h at room temperature. Of each hydrolysate, 100 µL was used to make a 10-fold dilution in ddH2O and the pH was neutralized using solid calcium carbonate. Solutions of 100 ppm D-glucose and D-xylose (Sigma) were used as standards. All sample were filtered through a 0.45 µm syringe filter before analysis. Glucose and xylose in the neutralized hydrolysates were measured using a high-performance anion-exchange chromatography Dionex ICS-6000 system (Thermo Scienctific). Monosaccharides were separated using ddH2O (A) and 200 mM sodium hydroxide (B) as the mobile phases at a flow rate of 0.4 mL/min through a Dionex CarboPac^TM^ PA20 analytical column (3 × 150 mm; Thermo Scientific) connected to a Dionex CarboPac^TM^ PA20 3 × 30 mm guard column (Thermo Scientific). Solvent B was used at 1.2% for the first 18 min and gradually increased to 50% by 20 min and run through 30 min. Solvent B was then reduced to 1.2% and run for 5 min (total run time was 35 min). Glucose and xylose were detected using a pulsed amperometric PAD; ICS-6000 detector (Thermo Scientific; Waltham, MA, USA) with a working gold electrode and a silver-silver chloride reference electrode at 2.0 μA.

#### Enzyme preparation

*T. reesei* Cellobiohydrolase I (Sigma catalog # E6412), henceforward Cel7A, was buffer exchanged into 50 mM borate buffer (pH 8.5) using Bio-Spin P-30 Bio-Gel spin-columns (Bio Rad). Enzyme concentration was determined using an extinction coefficient of 74,906 M^-1^cm^-1^ and 280 nm absorbance. Cel7A was biotinylated using EZ-Link NHS-LC-LC-Biotin (Thermo Scientific catalog # 21343), by combining the enzyme with biotin-NHS dissolved in anhydrous dimethylformamide (DMF) at a biotin:enzyme ratio of 10:1, and incubated for 4 h in the dark at room temperature. To remove unbound biotin, the enzyme+biotin mixture was buffer exchanged into 50 mM sodium acetate using Bio-Spin P-30 Bio-Gel columns. The biotinylated enzyme concentration was calculated again using absorbance measurements at 280 nm and the biotin concentration was determined using a Pierce Fluorescence Biotin Quantitation Kit (Thermo Scientific). Glycerol was added to biotinylated Cel7A to 10% v/v, and then 5 µL enzyme samples with a concentration of 6 µM were aliquoted, flash frozen in liquid nitrogen, and stored at −80°C until use.

#### Single-molecule microscopy

To prepare each flow cell, ∼10 µL of microfluidized cellulose/composite suspension was pipetted onto the surface of a glass slide. Two strips of double-sided tape were positioned on either side of the sample and a 18 x 18 mm glass cover slip was placed on top of the tape to create a flow cell (∼30 µL volume). The slides were inverted and placed into an oven at 60°C for 30 min to allow the cellulose solution to dry, leaving the cellulose/composite immobilized on the surface of the cover slip. A solution of 40-fold dilution of TetraSpeck beads (Thermo Scientific catalog # T7280) in 50 mM sodium acetate pH 5, used as fiduciary markers, was then injected into the flow cell and incubated for 5 min to allow the beads to bind to the cover slip surface. To prevent nonspecific binding of Cel7A to the glass surface, three washes of 1 mg/mL BSA in 50 mM sodium acetate, pH 5 were performed for 3 min each. Biotinylated Cel7A was diluted once in 50 mM sodium acetate, pH 5, to a concentration of 50 nM and then mixed with 20 nM (Thermo Scientific) Qdot-labeled Cel7A in 50 mM sodium acetate, pH 5.0, containing 5 mM dithiothreitol (DTT) to stabilize fluorescence. The final Qdot-labeled Cel7A working mixture contained 3 nM biotinylated Cel7A and 2 nM Qdot 655 was incubated for 15 minutes. The labeled enzyme solution was then injected into the flow cell. Single-molecule imaging was performed using total internal reflection fluorescence microscopy (TIRFM) with an excitation laser of 488 nm at 30 mW power to illuminate both the TetraSpeck beads on the surface and the enzyme-linked Qdots^19^. Substrate was imaged by interference reflectance microscopy (IRM) with a white light LED. Images were acquired at a rate of 1 frame/s for a total of 1,000 frames. The imaged field of view was 79.2 µm x 79.2 µm with a pixel size of 73 nm. To maintain constant focus during image acquisition, a quadrant photodiode (QPD) sensor connected to the microscope stage was used for real-time correction of z drift.

The acquired time-lapse stacks were analyzed using Fluorescence Image Evaluation Software for Tracking and Analysis (FIESTA) software ^20^. By fitting a two-dimensional Gaussian curve to the point-spread functions of imaged puncta (TetraSpeck beads and Qdots), FIESTA calculates the trajectories of labeled Cel7A molecules, and using the immobile TetraSpeck beads, also computes appropriate corrections for any XY drift during image acquisition. The traces generated using FIESTA were then further analyzed using a custom MATLAB-based pipeline^7,19^.

#### SEM and fibril measurements

To prepare samples for SEM imaging, 10 µL of substrate suspensions were pipetted onto filter membrane disks (Millipore, 0.2 µm). A series of 5 min ethanol washes followed, starting with 25% (v/v) ethanol followed by 50 %, 60%, 70%, 85%, 95%, and 100% ethanol. Critical point drying was performed using an EM CPD300 critical point dryer (Leica). Following drying, membranes with samples were mounted onto aluminum stubs using carbon tape and sputter coated with a 5 nm iridium coat using an EM ACE200 sputter coater (Leica). Samples were imaged with a SIGMA VP-FESEM (Zeiss) utilizing the secondary electron detector.

Fibril width measurements were performed manually on micrographs at 100,000 x magnification using ImageJ. At least 100 fibrils form 10 different micrographs for each substrate were measured.

#### Statistical analysis

All plots and statistical analysis were made using GraphPad Prism 10 software.

## Results and Discussion

To study the effects of xylan on cellulose deconstruction by Cel7A, we first synthesized cellulose-xylan composites to use as substrates. These composites were produced by growing *Gluconacetobacter hansenii*, a cellulose-synthesizing bacterium, in media supplemented with increasing concentrations of purified beechwood xylan. Cultures containing 0.05 µg/mL, 1 µg/mL, 200 µg/ml, or 1000 µg/mL xylan were used, and the resulting composites were imaged with fluorescence microscopy to monitor for the incorporation of xylan into the cellulose mesh. Staining with S4B for cellulose, in combination with anti-xylan immunofluorescence using LM11 antibodies, suggested a minute presence of xylan in the lowest, 0.05 µg/mL xylan concentration (Fig. 1D-F). More prominent xylan immunolabeling was evident in the composite grown with 1 µg/mL xylan (Fig. 1G-I), and its coverage of the S4B signal increased with the increase in media xylan concentration (Fig. 1J-O). A colocalization analysis using the S4B and xylan immunolabeling signals showed that the Pearson’s correlation coefficients (*r*_p_) increased together with the increase in media xylan concentrations up to the 200 µg/mL concentration. The 1000 µg/mL xylan substrate showed a slight reduction in *r*_p_ (Fig. 1P), possibly due to excess xylan that was separated from the composite. Evidence for such xylan that is not associated with cellulose staining can be seen in Fig. 2M-O, red arrowheads.

**Figure 1.**
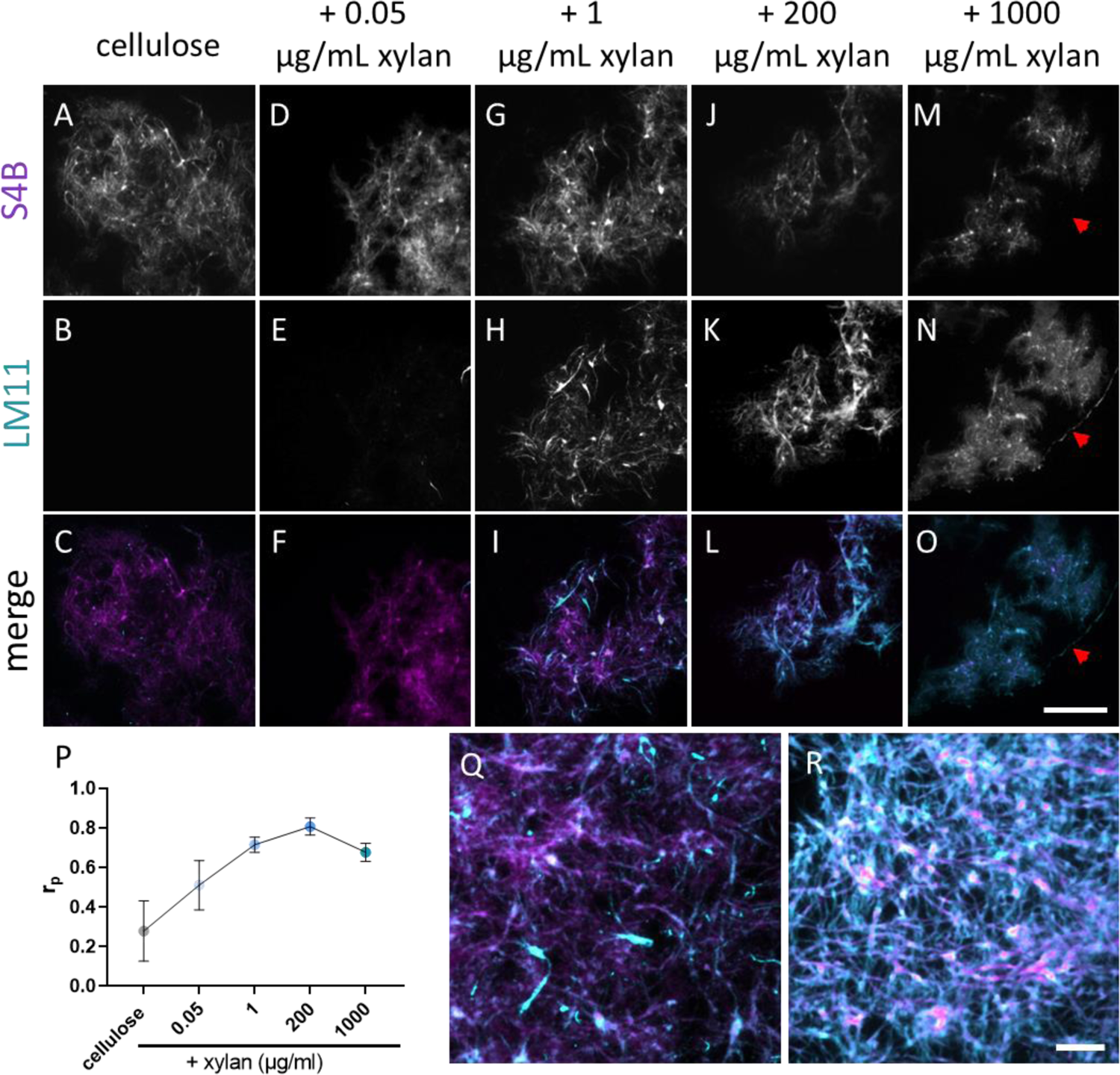
Incorporation of purified beechwood xylan into bacterial cellulose based composites. Bacterial cellulose staining using S4B (top row) and xylan immunofluorescence using LM11 antibodies and Alexa488-labeled secondary antibody (middle row). Merged images show the S4B signal in magenta and the Alexa488 signal in cyan (bottom row). P, *r*_p_ of colocalization analysis of cellulose and xylan signals in the different substrates. Q and R, representative merged images of 1 µg/mL and 200 µg/mL xylan samples, respectively. At least seven micrographs per sample were used to calculate *r*_p_. Scale bar in panel O, common to panels A-O, represents 50 µm. Scale bar in R, common to panels Q and R, represents 10 µm.

**Figure 2.**
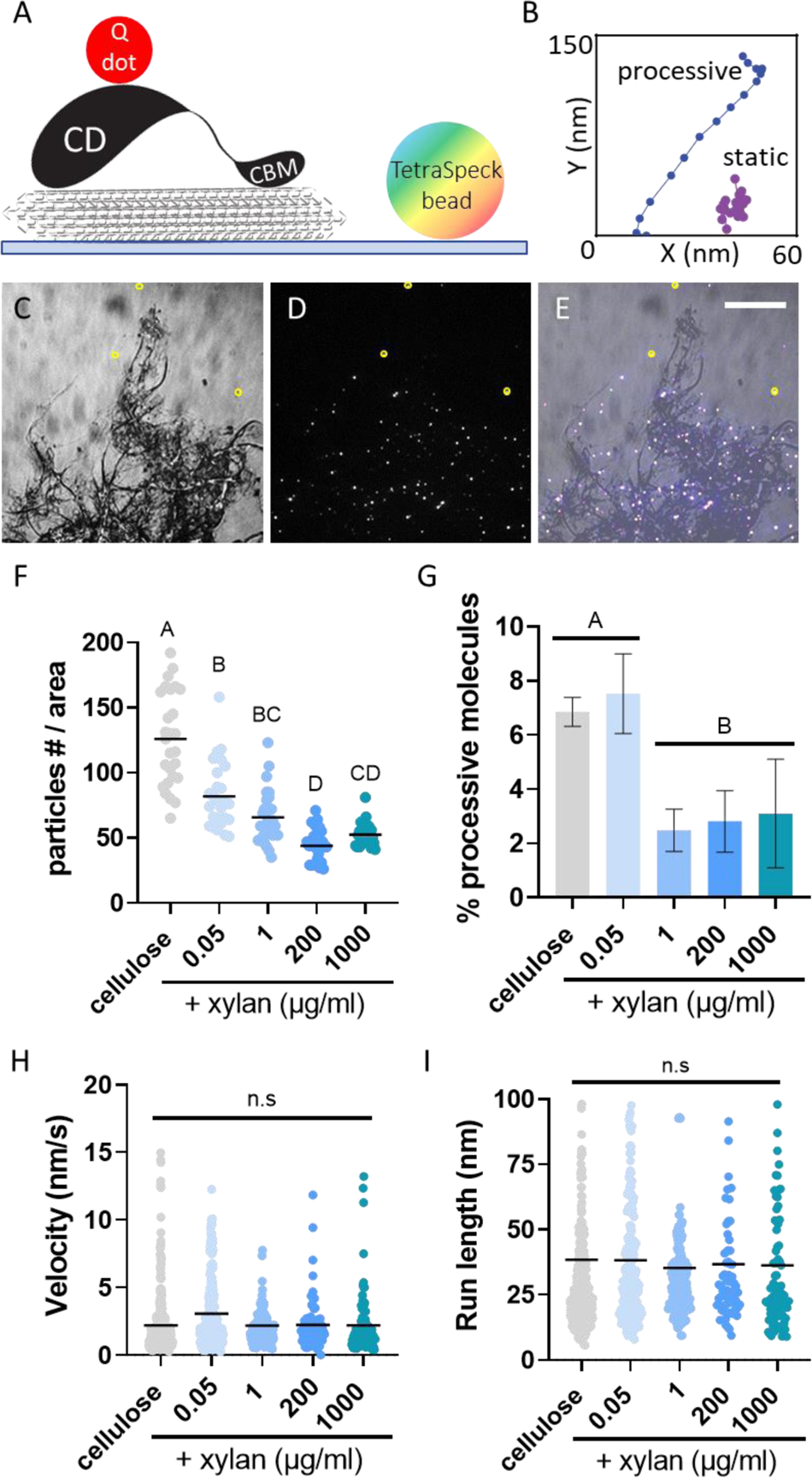
Single-molecule imaging of Qdot-labeled Cel7A. A, a cartoon of the experimental setup, showing a Q-dot labeled Cel7aA molecule on surface immobilized cellulose (seen in gray). A TetraSpeack bead, used as fiduciary marker, is immobile on the glass surface. B, representative displacement plots of a static particle (magenta) and a particle exhibiting processive movement (blue). C, IRM image of bacterial cellulose used as substrate for single-molecule experiments. D, TIRFM image of the sampe field of view seen in panel C taken at 90 sec after introducing labeled Cle7A into the cellulose containing flow cell. Labeled enzyme molecules, seen as white dots, populate the cellulose-containing area of the flow cell. E, merged image combining panels C and D. Three TetraSpeck beads are immobile on the glass surface and are circled in yellow in panels C-E. Scale bar in panel E, common to panels C-E, represents 20 µm. F, G, H and I, mean of number of bound particles at steady-state, mean proportion of processive particles, mean velocity and mean run length, respectively. Binding was calculated from three images from three different experiments for each substrate. The proportion of processive molecules was calculated from at least three experiments. The velocity and run length were calculated from at least 100 processive particles from three different experiments. A one-way ANOVA and Tukey’s multiple comparison test was used to compare the different substrate. Different letters indicate statistical difference (p<0.05).

To assess whether xylan interferes with the binding of Cel7A to cellulose, we performed single-molecule binding assays by allowing Qdot-labeled Cel7A enzyme to reach steady-state binding equilibrium with the different substrates over 5 min, then recorded the number of bound molecules per area. The number of bound enzymes was inversely correlated with the xylan content in the substrate: even in the lowest xylan-containing composite, made with 0.05 µg/mL xylan, the number of bound enzymes was reduced by ∼50% as compared to the cellulose-only control, and a maximum reduction of ∼65% was reached in the 200 µg/mL xylan composite (Fig. 2G). As we did not notice any preferential binding at specific locations in the composites, these data indicate that Cel7A molecules do not favorably bind to xylan when it is present, but instead are inhibited from binding to cellulose.

We then analyzed the effects of xylan on the motility of the Cel7A molecules that did bind to the substrate by imaging and tracking the binding and processive movements of Qdot-labeled enzyme on the different substrates. By analyzing traces of the labeled molecules, we calculated the proportion of processively moving enzymes, their velocity, and the length of their processive runs. On cellulose alone and the 0.05 µg/mL xylan composite, ∼7% of bound Cel7A molecules exhibited processive movement, in accordance with our previous work^7^. All other xylan-containing substrates showed a significantly reduced proportion of moving Cel7A particles with 2 – 4% of imaged enzyme molecules displaying at least one processive run (Fig. 2H). The velocities and run lengths of Cel7A molecules were similar between all tested substrates with means of 2 – 3 nm/s and ∼35 nm, respectively; both of these values consistent with previously published data (Fig. 4I-J)^7^. To test whether the source of xylan might affect these findings, we also used composites assembled with 1 µg/mL purified oat xylan or wheat arabinoxylan, that have differences in composition and structure from beechwood xylan. Similar results, with some reduction in run lengths, were obtained (Fig S1), indicating that the ability of different types of xylans to inhibit Cel7A binding and motility does not depend on their specific chemical compositions or sidechain configurations.

To more precisely quantify the extent of xylan inclusion in the different substrates, we performed a monosaccharide analysis of acid-digested substrates using high performance liquid chromatography (HPLC). As expected, a glucose peak with an elution time of 8-8.5 min was evident in all tested samples (Fig. 3A). A xylose peak, with an elution time of 9-10 min, could be seen in composites grown with 200 and 1000 µg/mL xylan. The ratios of the xylose and glucose peak areas were 0.6717±0.01 (xyl:glu) and 1.224±0.01 in the 200 µg/mL and in the 1000 µg/mL xylan-containing substrates, respectively (Fig. 3B). However, in the 0.05 µg/mL and 1 µg/mL xylan samples, xylose concentrations were below the detection limit (Fig. 3A).

**Figure 3.**
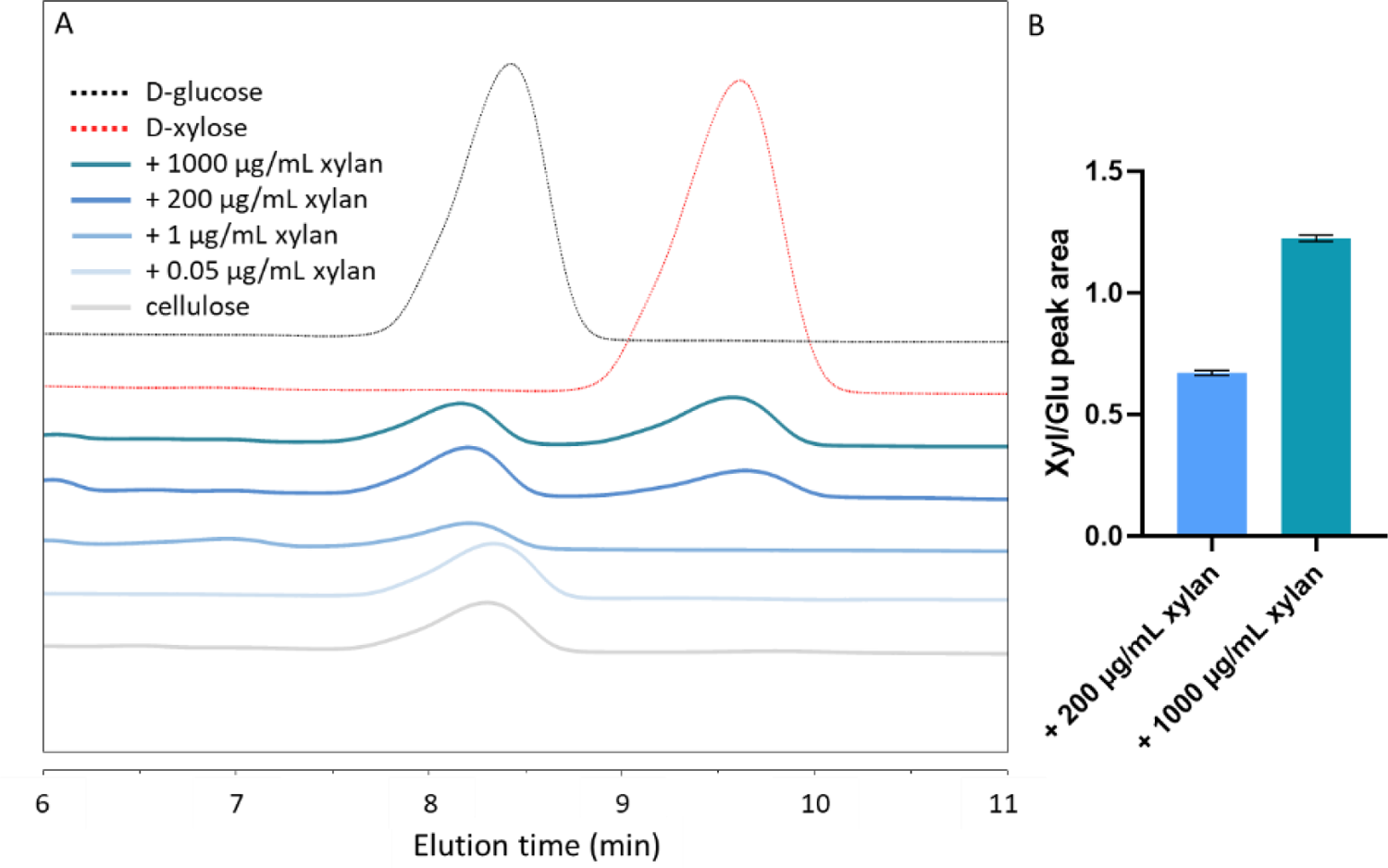
Monosaccharide analysis of acid digested composites. A, HPLC measurements of xylose and glucose content in hydrolyzed samples. B, Peak areas were used to evaluate the ratios between xylose and glucose. Chromatograms are representative of two experiments using separately hydrolyzed substrates.

To gain a better understanding of the mesoscale distributions of xylan within the composites and how they relate to our single-molecule results, we examined the different substrates using scanning electron microscopy (SEM). All substrates showed the typical fibrillar lattice of bacterial cellulose (Fig. 4). Some non-fibrillar components could occasionally be found in the substrate grown with 1 µg/mL xylan. These nebulous structures partially covered and bridged adjacent cellulose fibrils (Fig. 4C, red arrowheads), but were irregularly dispersed in the sample. More noticeable structural differences could be seen in the 200 µg/mL xylan substrate, where patches of seemingly coagulated cellulose fibrils were frequently found throughout the sample (Fig. 4D). These patches appeared to bridge numerous fibrils and in certain cases created continuous sheet-like structures (Fig. 4D, red arrowheads). In the sample grown with the highest xylan concentration tested, 1000 µg/mL, complete areas of the cellulose mesh appeared to be cemented to the extent that, in some cases, cellulose fibrils were entirely covered and were difficult to identify (Fig. 4E, red arrowheads). Taken together, these results suggest that at low concentrations, xylan incorporates into the composite in an unstructured association with individual cellulose fibrils, whereas when higher concentrations of xylan are supplied, it accumulates into larger patches covering tens or hundreds of fibrils. However, it should be noted that all xylan-containing composites exhibited highly heterogeneous morphologies.

**Figure 4.**
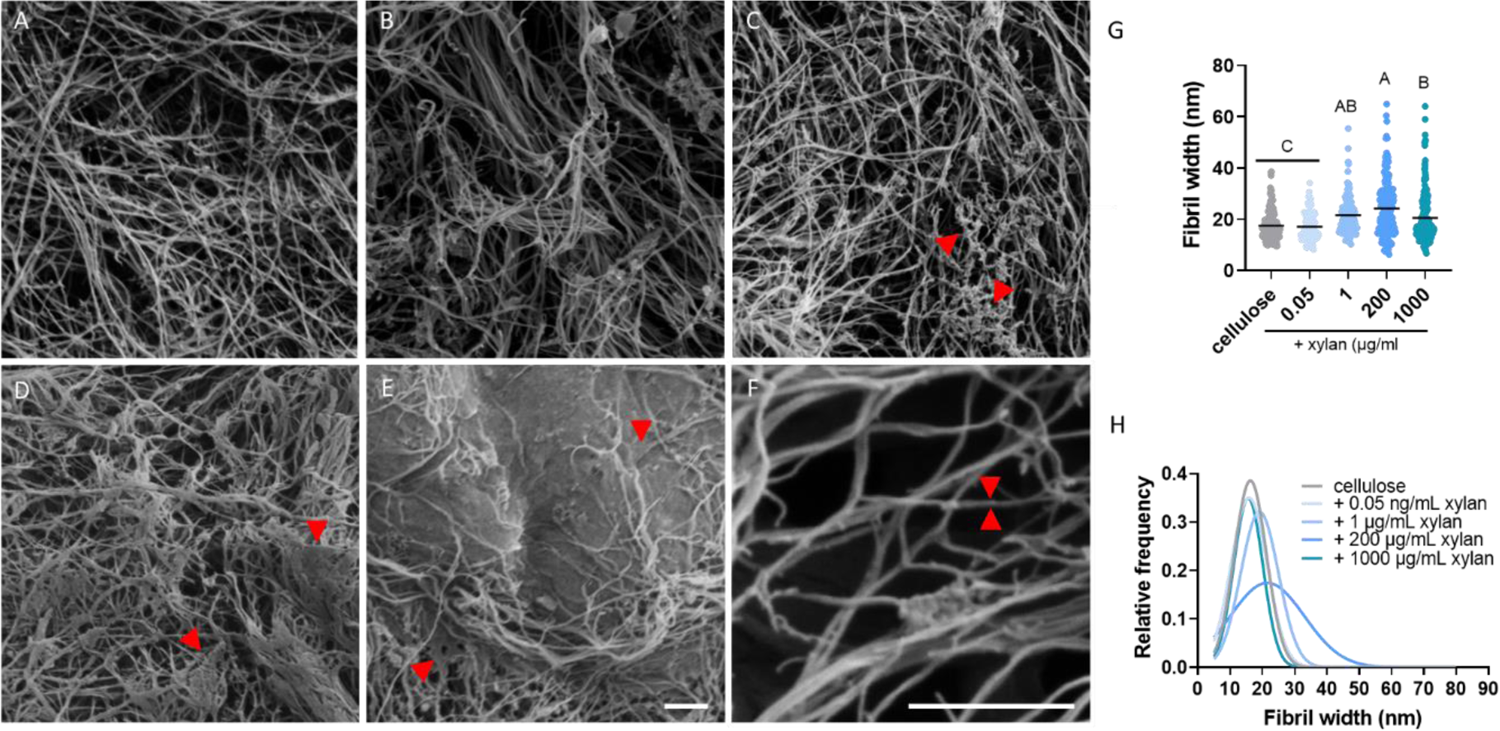
SEM micrographs of bacterial cellulose and xylan-containing composites. A, bacterial cellulose with no added xylan. B, C and D, composites containing 0.05 µg/mL, 1, 200 and 1000 µg/mL xylan, respectively. F, micrograph used to measure single fibril width, with a representative fibril highlighted by two red arrowheads. G, mean fibril width in the different substrates. H, Gaussian fit of fibrils width distribution of each of the substrates detailed histogram for panel H can be found in figure S2. At least 100 fibrils from 10 different micrographs for each substrate were measured. Scale bar in E common to panels A-E. Scale bars represent 300 nm. A one-way ANOVA and Tukey’s multiple comparison test was used to compare the fibrils width in different substrate. Different letters indicate statistical difference (p<0.05).

Considering the reduction in enzyme binding observed in all the xylan-containing substrates (Fig. 2G), and the reduced proportion of processive enzymes in the 1, 200, and 1000 µg/mL substrates (Fig. 2H), we hypothesized that xylan also accumulates as a thin coating surrounding cellulose fibrils. Such a coating might prevent Cel7A from binding to its substrate, prevent successful extraction of a glucose chain from the cellulose surface^6,9,21^, or block the movement of enzyme molecules that do engage with the cellulose. We therefore compared the widths of individual fibrils in the different substrates, measuring the thinnest fibrils present in each micrograph. The mean fibril width in the cellulose-only control was 17.5 nm, significantly lower than the 21.5 nm, 24.2 nm, and 20.5 nm measured in samples grown with 1, 200, and 1000 µg/mL xylan, respectively. However, fibril width did not significantly increase relative to the cellulose-only control in the 0.05 µg/mL samples (Fig. 4G). Comparing the distribution of fibril widths measured from the different substrates, all xylan-containing substrates had a larger proportion of thicker fibrils (Fig. 4G-H). These data can be interpreted in at least two ways: first, xylan coatings might thicken individual cellulose fibers when the cellulose is synthesized in the presence of xylan, and second, the xylan might adhere cellulose fibrils together, causing them to coalesce into thicker bundles.

The efficiency of producing biofuel and other bioproducts from lignocellulosic biomass is limited by the productivity of cellulolytic enzymes. Specifically, non-cellulosic cell wall components, such as xylan, are known to inhibit the activity of cellulase enzymes, thereby increasing the recalcitrance of plant biomass. Here, we aim to shed light on the mechanisms underlying this inhibition.

By imaging Qdot-labeled Cel7A by TIRFM, we measured the dynamics of the enzyme at single-molecule resolution. Our results show a considerable reduction in enzyme binding in all xylan-containing substrates when compared to the cellulose-only control. Notably, this reduction took place in the composites grown with 0.05 µg/mL and 1 µg/mL xylan, where xylose concentrations were too low to detect by our monosaccharide analysis (Fig 3). However, a reduction in the proportion of processive enzyme molecules was evident in the 1 µg/mL sample, but not in the 0.05 µg/mL sample. This may hint at the different ways in which xylan incorporates into the cellulose network when supplied at different concentrations (Fig. 5). At 0.05 µg/ml, xylan might sparsely coat the cellulose such that some Cel7A binding sites become inaccessible for the enzyme, but not do so to the extent that it prevents the movements of Cel7A molecules that do bind. With increasing xylan concentrations, its coverage of the cellulose network also increases and the availability of binding sites further declines. A slight, nonsignificant, increase in binding seen in the 1000 µg/mL sample might be explained by phase separation and aggregation of xylan separately from the cellulose. At concentrations of 1 µg/mL and higher, in addition to blocking potential binding sites, dense xylan coating on the cellulose surface might prevent bound molecules from successfully complexing with a glucose chain or constitute xylan “roadblocks” that obstruct processive progress for complexed Cel7A molecules.

**Figure 5.**
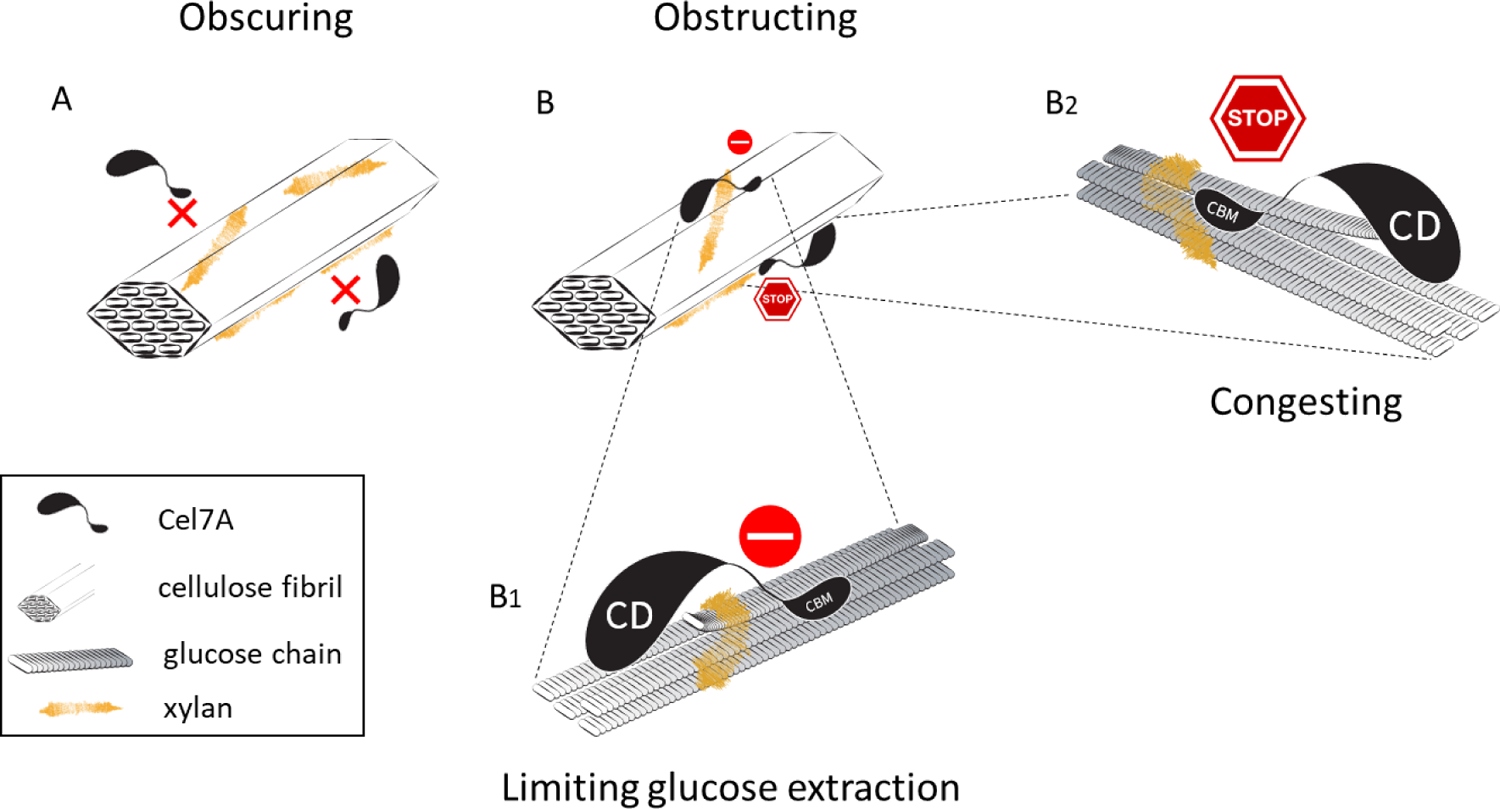
Mechanisms of xylan inhibition of Cel7A. The results of this study indicate that xylan interferes with Cel7A activity in two distinct ways. A, Obscuring-xylan leads to reduced numbers of binding sites accessible to the enzyme either through direct coverage of these sites, or indirectly by increasing fibril bundling, which reduces the effective surface area of cellulose. B, Obstructing-xylan prevents the processive movement of enzyme molecules that successfully bind to cellulose. We suggest two possible explanation for this obstruction: B1, inhibiting glucose extraction-xylan bound to cellulose stabilizes the surface glucose molecules and makes the threading of a glucose chain into the catalytic tunnel less favorable. This reduces the probability of effective enzyme-cellulose complexing and consequently the proportion of processively moving enzyme molecules. B2, congesting-xylan on the surface of cellulose physically blocks the movement of the enzyme. This can occur even when an enzyme molecule is successfully complexed, and after a processive run had begun.

To generate our synthetic xylan containing composites, we supplemented cultures of *Gluconacetobacter hansenii* with variable concentrations of purified beechwood xylan, and found that we can control the extent of xylan incorporated in the resulting pellicles. Our colocalization analysis, which was based on the degree of overlap between the S4B and xylan Alexa 488 signals, showed a maximal *r*_p_ mean value of 0.8 in the 200 µg/mL samples. The reduction in *r*_p_ found in the 1000 µg/mL sample relative to that value might result from xylan that is not associated with cellulose, possibly due to phase separation and xylan aggregation in solution, or the detaching of loosely bound xylan from the cellulose network. In either case, this finding indicates that by supplementing cultures with xylan at a concentration of 1000 µg/ml, we were reaching saturation of the composite. When this sample was dehydrated, xylan that aggregated in solution was likely deposited on the sample and underlying surfaces. This can be seen in our immunofluorescence results (Fig. 1M-O) and can also explain the patches of entirely covered fibrils found in SEM (Fig. 4E).

Studies of xylan biosynthesis and its interactions with cellulose in plants have demonstrated its importance for proper cell wall development^22–24^. In *Arabidopsis thaliana*, multiple mutants defective in xylan biosynthesis are known. Many of these are *irregular xylem* (*irx*) mutants, which are characterized by formation of irregularly shaped xylem vessels. While these mutations impact different steps of the xylan biosynthesis pathway, many phenotypes include defective coalescence and orientation of cellulose microfibrils that lead to compromised mechanical integrity of the cell walls, and in some cases to considerable dwarfism of the entire plant^22,25^. Evidence for the effects of xylan abundance and binding availability during cellulose synthesis on the bundling of cellulose^3,23^ can also be found in our *in-vitro* generated composites. We observed large-scale aggregates, presumably of xylan, at higher concentrations of xylan addition. Additionally, although our SEM imaging did not show obvious effects of xylan supplementation on the morphology of the thinnest fibrils evident in the micrographs, we measured an increase in the mean width of those fibrils. This can be explained by I) the incorporation of xylan in between glucose chains and sub-elementary fibrils, II) the covering of cellulose fibrils by a thin layer of xylan, and/or III) the formation of higher-order cellulose bundles. The expanded distribution of fibril widths observed in our xylan-containing composites (Fig. 4G-H) supports the latter explanation. Additionally, the reduction in bound enzyme molecules found in all the xylan-containing substrates suggests that at least part of the cellulose network becomes unavailable for enzyme binding, perhaps due to a covering of xylan that is undetectable by SEM but is sufficient to make potential binding sites inaccessible.

Molecular dynamics simulations of xylan and cellulose interactions suggest that short xylan oligomers tend to migrate towards the hydrophobic planes of cellulose fibrils and become stabilized on them^1^. Similar simulations found that the diffusion of the Cel7A CBM towards and on hydrophobic surfaces of cellulose is thermodynamically favorable^26^. This favored localization of both xylan and enzyme to the hydrophobic surfaces might explain why even residual amounts of xylan are sufficient to significantly affect Cel7A binding and motility. In this case, complete coating of cellulose fibrils is not necessary to effectively inhibit the enzyme binding. Moreover, interactions between and bundling of neighboring fibrils will likely involve the hydrophobic surfaces on which xylan accumulates and will lead to further reduction in accessible binding sites for Cel7A. This can explain the concentration dependent effect of xylan supplementation on enzyme binding that we observe (Fig. 2G). Xylan bound to the hydrophilic surfaces of cellulose assumes a twofold screw configuration and is bound by complex hydrogen bonding. In this conformation xylan is seen as an extension of the crystalline structure of the cellulose fibril^27^, and is suggested to stabilize surface glucose residues, making them less mobile^1^. This might also be expected to interfere with extraction of glucose chains from the cellulose surface and limit the complexation of Cel7A with its substrate.

A commonly suggested inhibitory effect of xylan is that it drives off-target enzyme binding. According to this approach, xylan acts as a sink that depletes Cel7A molecules, consequently reducing enzyme efficiency^28,29^. No evidence for such an inhibitory mechanism was found under our experimental conditions. However, it is possible that nonspecific binding of cellulases and other wall-degrading enzymes to xylan becomes more relevant under industrial conditions, where elevated temperatures and high enzyme loadings are applied.

The fact that xylan inhibits Cel7A when present at almost undetectable amounts has important implications for biomass saccharification at industrial scales, where xylan content remains substantial even after pretreatments. In corn stover, hemicellulose was found to make up 6%-23% (w/w) of pretreated biomass^30^, and comparable proportions were reported for sugarcane bagasse^31^. At these levels of xylan, which correspond roughly to the addition of 1-200 µg/mL of xylan in our synthetic composites, Cel7A efficiency is expected be hindered by two mechanisms: inhibition would arise from both reduced binding and the impediment of bound enzyme molecules (Fig. 5).

## Conclusion

By tracking *Tr*Cel7A molecules at single-molecule resolution on cellulose alone and cellulose-xylan composites, we were able to gain new insights into how xylan affects the dynamics of this economically important enzyme. Our results show a double effect of xylan on Cel7A efficiency: I) by reducing its binding to cellulose, most likely by masking potential binding sites on the cellulose surface, and II) by preventing bound enzymes form processively advancing on the substrate, possibly by hindering extraction of glucose chains form the cellulose crystal or by causing “road blocks” that obstruct processivity. Our data suggest that even small amounts of xylan coat cellulose in a way that inhibits Cel7A binding, and that incorporation of additional xylan promotes the bundling of cellulose fibrils, possibly at multiple scales, thus leading to the formation of higher-order cellulose ribbons and sheets. These results demonstrate the molecular impacts of retaining residual xylan during biomass processing and enzymatic saccharification, and stress the importance of targeting xylan separation and/or digestion during biomass processing as a strategy for improving the efficiency of biomass conversion into biofuels and bioproducts. To capture the effects of xylan on lignocellulose saccharification in native plant cell walls, our single-molecule tracking approach could be combined with plant-derived lignocellulose substrates to shed additional light on the potential and limitations of these enzymes and aid in developing superior enzymes and strategies for the effective use of biomass in the sustainable bioeconomy.

## Author Contributions

WH and CTA obtained funding, CTA and NZ designed experiments, NZ and AP performed experiments and analyzed data, NZ wrote the paper, and NZ, CTA, AP, DN, and WH edited the paper.

## Conflicts of Interest

The authors declare no conflicts of interest.

## Acknowledgments

This research was supported by the Department of Energy Office of Science grant number DE-SC0019065, the National Science Foundation grant number 2301377, and by BARD, the United States - Israel Binational Agricultural Research and Development Fund, Vaadia-BARD Postdoctoral Fellowship Award FI-622-2022.

## Supplementary

**Figure S1.**
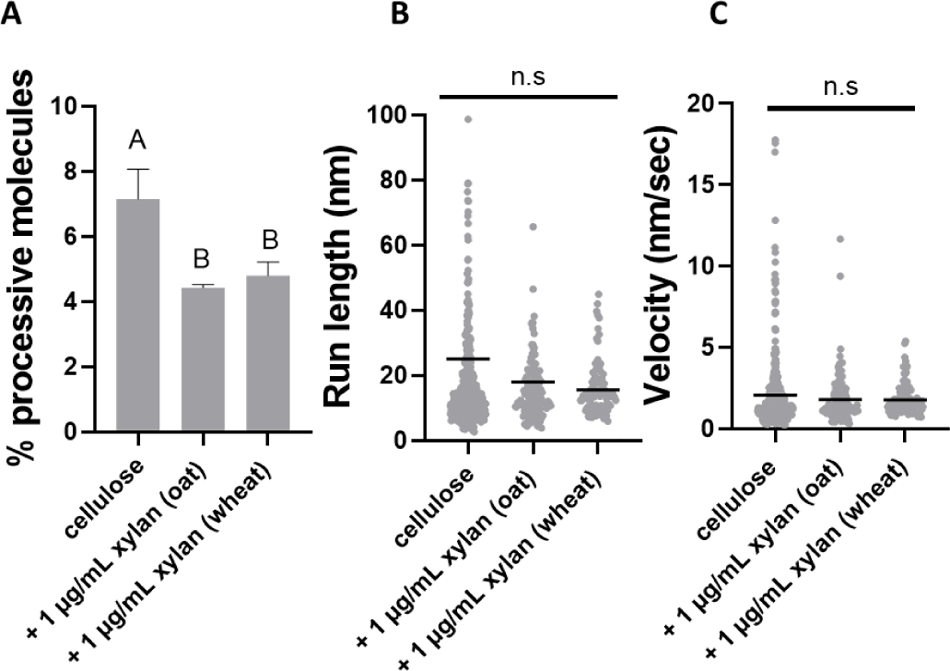
Single molecule dynamics of Cel7A on bacterial cellulose composites synthetized with 1 µg/mL purified oat xylan and wheat arabinoxylan. A, a reduction in percentage of processive molecules was similar to that found in beechwood xylan containing composites. B, some nonsignificant reduction in run length was observed in the composites as compared to the cellulose control. C, as with beechwood xylan, no significant differences were found in the velocities. Different letters indicate statistical difference (p<0.05).

**Figure S2.**
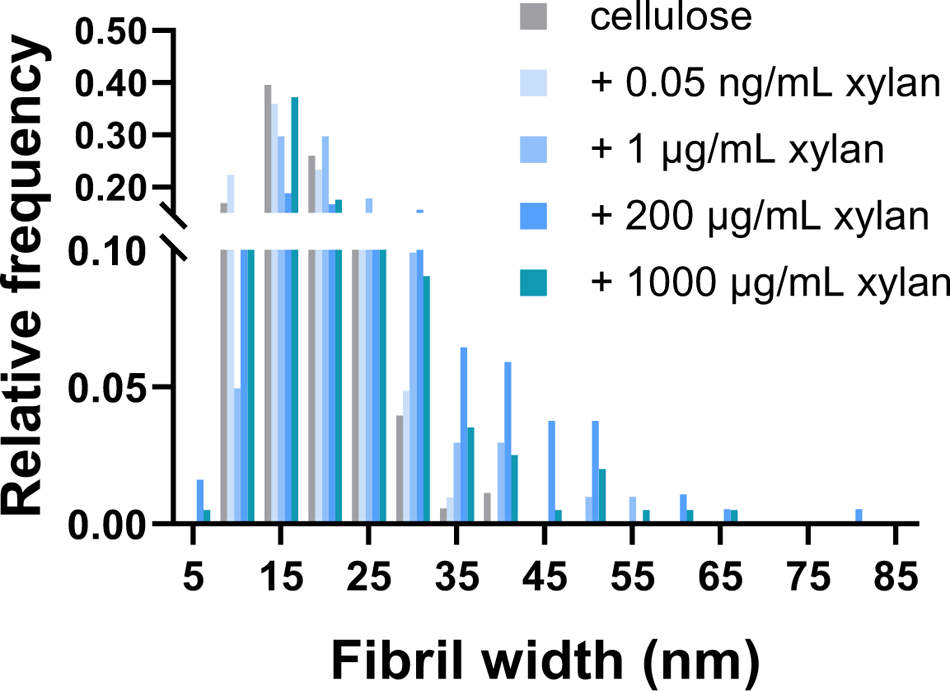
The distribution of fibril width of cellulose and all xylan-containing composites measured from SEM micrographs. This histogram was used to generate the fit curves shown in figure 4H.

## Notes

### Competing Interest Statement

The authors have declared no competing interest.

